# A single heterozygous mutation in *COG4* disrupts zebrafish early development via Wnt signaling

**DOI:** 10.1101/2021.05.23.443307

**Authors:** Zhi-Jie Xia, Xin-Xin I. Zeng, Mitali Tambe, Bobby G. Ng, P. Duc S. Dong, Hudson H. Freeze

**Author notes:** **Competing Interests**: The authors have declared that no competing interests exist.

## Abstract

Saul-Wilson syndrome (SWS) is a rare, skeletal dysplasia with progeroid appearance and primordial dwarfism. It is caused by a heterozygous, dominant variant (p.G516R) in COG4, a subunit of the Conserved Oligomeric Golgi (COG) complex involved in intracellular vesicular transport. Our previous work has shown the intracellular disturbances caused by this mutation; however, the pathological mechanism of SWS needs further investigation. We sought to understand the molecular mechanism of specific aspects of the SWS phenotype by analyzing SWS-derived fibroblasts and zebrafish embryos expressing this dominant variant. SWS fibroblasts accumulate glypicans, a group of heparan sulfate proteoglycans (HSPGs) critical for growth and bone development through multiple signaling pathways. Consistently, we find that glypicans are increased in embryos expressing the *COG4*^*p.G516R*^ variant. These animals show phenotypes consistent with convergent extension (CE) defects during gastrulation, shortened body length, and malformed jaw cartilage chondrocyte intercalation at larval stages. Since non-canonical Wnt signaling was shown in zebrafish to be related to the regulation of these processes by Glypican 4, we assessed *wnt* levels and found a selective increase of *wnt4* transcripts in the presence of COG4^p.G516R^. Moreover, overexpression of *wnt4* mRNA phenocopies these developmental defects. LGK974, an inhibitor of Wnt signaling corrects the shortened body length at low concentrations but amplifies it at slightly higher concentrations. WNT4 and the non-canonical Wnt signaling component phospho-JNK are also elevated in cultured SWS-derived fibroblasts. Similar results from SWS cell lines and zebrafish point to altered non-canonical Wnt signaling as one possible mechanism underlying SWS pathology.

## Introduction

Saul-Wilson syndrome (SWS) is a rare skeletal dysplasia characterized by profound short stature and distinctive craniofacial features such as prominent forehead, prominent eyes and micrognathia (1, 2). Recently, we defined a specific heterozygous COG4 substitution (p.G516R) as the molecular basis of this rare form of primordial dwarfism (3). COG4 is one of the eight subunits of the conserved oligomeric Golgi (COG) complex regulating protein trafficking and Golgi homeostasis (4). Biallelic pathogenic variants in *COG4* and other COG subunits cause multiple human congenital disorders of glycosylation (CDGs) (5). COG4-CDG individuals have a very severe, usually lethal, phenotype with dysmorphia, neurological and intellectual disabilities and altered N-glycosylation with an almost total loss of COG4 (6, 7). However, SWS subjects show very different features from COG4-CDG individuals, and their N-glycans, intellectual and neurological features appear normal (3). At the cellular level, the COG4-p.G516R variant disrupted protein trafficking by accelerating Brefeldin-A (BFA)-induced retrograde transport and delaying anterograde transport, causing the collapse of the Golgi stacks. This interrupted bidirectional trafficking between the ER and the Golgi altered decorin, a proteoglycan (3), indicating modified proteoglycans may be involved in the pathogenesis of SWS.

Proteoglycans play critical roles in multiple cell processes at the cellular, tissue, and organismal level, and their deficiencies cause bone and connective tissue disorders (8, 9). Several proteoglycan deficiencies have been studied in zebrafish, a powerful vertebrate model for studying CDGs and skeletal disorders, with some showing shortened body axis (10-14). Besides a body axis defect, *decorin (dcn)* mutant zebrafish showed relatively severe defects in body curvature associated with a curved or not fully extended tail (11). Defects in glypicans, a group of heparan sulfate proteoglycans (HSPGs), can cause abnormal skull and skeletal dysplasia in both humans (Simpson-Golabi-Behmel syndrome, Glypican 3, GPC3; Keipert Syndrome, Glypican 4, GPC4) and zebrafish (Knypek, Gpc4) (15, 16). Interestingly, studies of *knypek* (*kny, gpc4*) mutant zebrafish demonstrated that Gpc4 deficiency causes chondrocyte stacking and intercalation defects in Meckel’s cartilage, which is not seen in *dcn* or *chondroitin sulfate proteoglycan 4* (*cspg4*) deficient zebrafish (11, 12, 17, 18). *kny* adult zebrafish also show craniofacial defects including a smaller head, domed skull and shorter jawbones, reminiscent of some of the clinical features of SWS individuals (16, 17). Studies also found that optimized expression of *gpc4* can suppress the defects caused by Wnt11f2 (formerly known as Wnt11/Silberblick (19)) deficiency, indicating the role of Gpc4 in the Wnt signaling pathway, probably as a Wnt coreceptor (16). Worth noting, both absence and high expression of *gpc4* leads to lost ability to suppress Wnt11f2 deficiency (17). Presumably, abnormalities arise when either these receptors or ligands are outside the optimal ratio. Taken together, the similarity between *kny* mutant zebrafish craniofacial defects and some of the SWS individuals’ clinical features encouraged us to use zebrafish as a vertebrate model to explore the underlying pathological mechanism of SWS.

The zebrafish Cog4 is 72% identical to human COG4 and the amino acid corresponding to the SWS mutation site is conserved across multiple vertebrate species (www.uniprot.org). Zebrafish that lack the Cog4 protein show phenotypes consistent with the clinical features of COG4-CDG individuals, including defective synthesis of N- and O-linked glycans, and decreased glycosphingolipid complexity (20). SWS individuals show very different features compared to COG4-CDG individuals, and SWS cells show accelerated BFA-induced retrograde trafficking opposite to COG4-CDG cells. Considering these facts, a zebrafish model for the SWS-specific variant is highly desired to investigate phenotypic features and the possible pathogenesis of this heterozygous mutation in *COG4*.

In this study, we utilize SWS-derived fibroblasts and a zebrafish system to test a specific heterozygous COG4 substitution (p.G516R) which is causal for SWS. We assessed a broader category of proteoglycans and found a consistent increase of glypican level in SWS-derived fibroblasts and zebrafish expressing human *COG4*^*p.G516R*^ variant. Further studies on non-canonical Wnts revealed that the presence of COG4^p.G516R^ specifically elevated *wnt4* transcript, but not *wnt5a, wnt5b*, or *wnt11f2*. Overexpression of *wnt4* phenocopies the *COG4*^*p.G516R*^ zebrafish, and the Wnt inhibitor LGK974 suppresses the defects caused by the expression of *COG4*^*p.G516R*^. These findings suggest that disrupted Wnt signaling is one possible mechanism underlying the pathogenesis of SWS.

## Results

### Fibroblasts from Saul-Wilson syndrome individuals accumulate glypicans on the cell surface

As one of the major components of extracellular matrix (ECM) and cell membrane proteins, proteoglycans comprise a large, heterogeneous group including HSPGs and chondroitin sulfate proteoglycans (CSPGs). Decorin is a predominant proteoglycan in human skin, covalently linked with one glycosaminoglycan chain (GAG) which requires normal function of the Golgi for its posttranslational modification. Although *dcn* mutant zebrafish does not show much clinical similarity to SWS individuals, nor is *Dcn* mRNA abundance changed in SWS-derived cells (unpublished data), abnormal decorin encouraged us to study other proteoglycans in SWS cells. As a first step in determining which proteoglycan may change, we analyzed the core proteins of HSPGs using an antibody (3G10) against HS-stubs that appear only after heparinase III digestion. As shown in Figure 1A, glypicans, syndecan 1 and syndecan 2 were the most prevalent cell-surface HSPGs in dermal fibroblasts. Compared to syndecans, glypicans showed consistent increases in all three SWS cell lines with an average 2-fold elevation (Figure 1B). We performed qPCR to assess whether transcript abundance is also altered. There are 6 glypicans in humans and, among those, glypican 1, 4 and 6 are present in multiple tissues, while other glypicans 3 and 5 are restricted to ovary and brain. Glypican 2 is specifically expressed in nervous system during embryonic development (21, 22). As seen in Figure 1C, glypican 1, 4, 5, and 6 were detectable in dermal fibroblasts; however, we did not observe a significant change in any of their transcript abundance (Figure 1D). We hypothesize this increase of glypicans results from SWS COG4-dependent abnormal trafficking, instead of transcriptional regulation. A similar strategy was applied to study CSPGs. Chondroitinase ABC was used to digest CSPGs followed by immunoblotting against the ΔDi-6S. There was a general decrease of CSPG core proteins in 3 out of 4 SWS-derived cell lines (Figure S1). Considering glypicans show the most prominent difference between controls and SWS-derived cells, and the phenotypes of *gpc4* mutant zebrafish, we focused on glypicans in our study.

**Figure 1.**
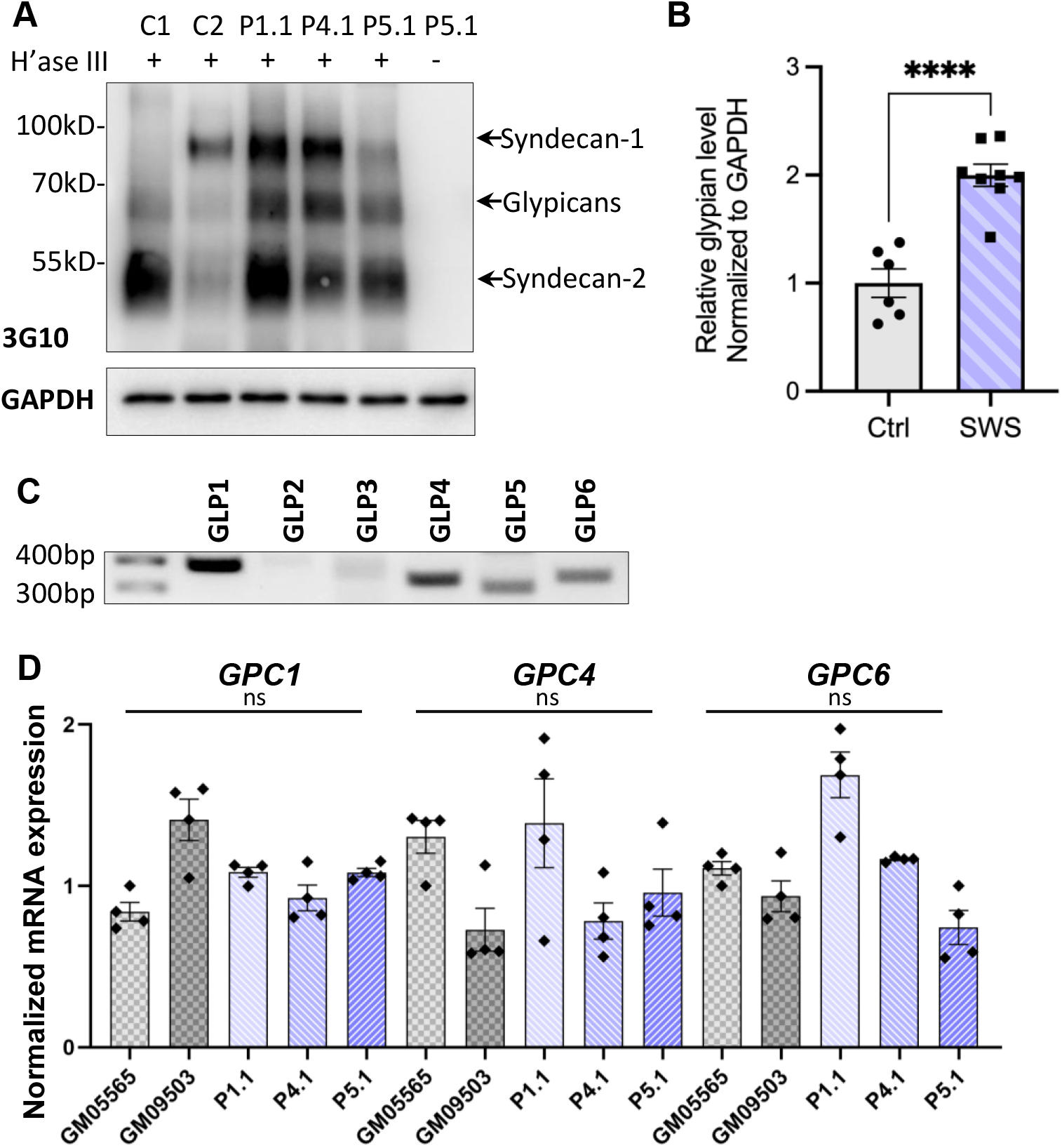
SWS-derived fibroblasts show altered HSPGs and glypicans after heparinase III digestion. (A) Western blotting of ΔHS-stub using 3G10 antibody following heparinase III digestion of three SWS-derived fibroblasts and two control fibroblasts. C1 and C2 are control fibroblasts. C1, GM08429; C2, GM08680. P1.1, P4.1 and P5.1 are SWS-derived fibroblasts. (B) Quantitation assay of glypican band density in (A) and two other replicates. The data are presented as mean ± SEM. Unpaired two-tailed *t*-test was used. ****, p<0.0001. (C) Agarose gel of six human glypicans after reverse transcript PCR. (D) qPCR of three dominant glypicans in SWS-derived fibroblasts. Relative glypican level was normalized to GAPDH. The graphs represent the 2^−ΔΔCt^ values. Experiments were performed in triplicates with similar results.

### Expression of human *COG4*^*p.G516R*^ in zebrafish increases glypican proteins

To study the impact of the SWS variant on skeletal development, we used zebrafish as a vertebrate model. Since SWS is a dominant disorder, we overexpressed the human SWS allele in developing zebrafish embryos. Human *COG4*^*WT*^ and *COG4*^*p.G516R*^ mRNA or DNA constructs, as shown in Figure 2A, were injected into one-cell stage embryos. The presence of human COG4 in zebrafish was confirmed by an antibody specifically recognizing human COG4 protein (Figure 2B-2C). Maintaining endogenous protein level of COG4 is important for its cellular function based on *in vitro* cell studies (23). Lacking a zebrafish Cog4 antibody makes it impossible to determine the relative expression level of *COG4*^*p.G516R*^, therefore we include *COG4*^*WT*^ as an injected control to ensure reasonable expression of *COG4*^*p.G516R*^ in zebrafish.

**Figure 2.**
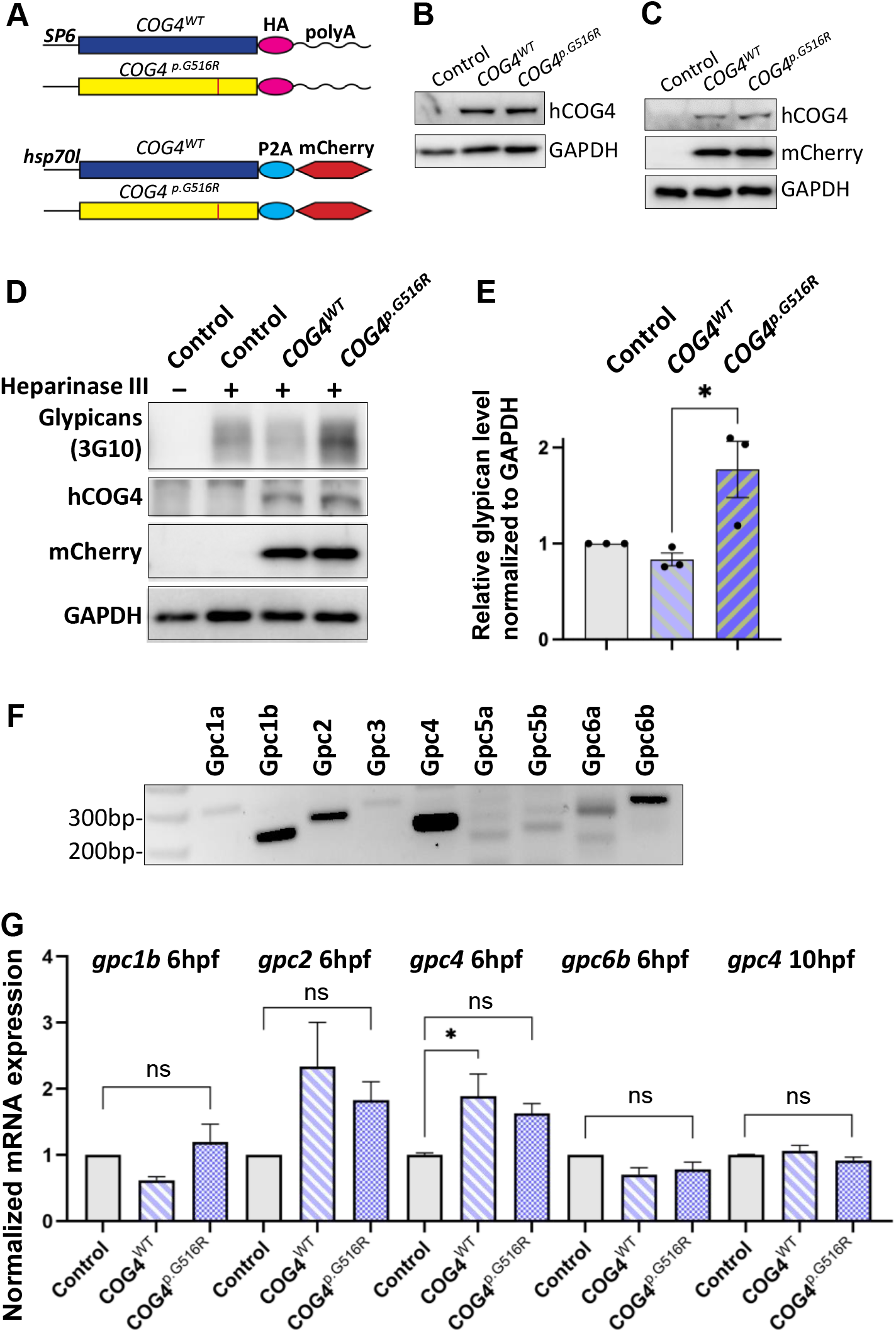
Expression of human *COG4*^*p.G516R*^ in zebrafish increases the protein level of glypicans. (A-C) Expression of human *COG4*^*WT*^ and *COG4*^*p.G156R*^ in zebrafish after mRNA or DNA injection. (A) The scheme of *COG4* constructs for *in vitro* transcription (top) and DNA injection (bottom). (B) Western blot at 24 hpf to detect the presence of COG4 after mRNA injection. (C) Western blot at 48 hpf to detect COG4 after DNA injection, heat shock was performed at 24 hpf for 2 hours at 38°C degree. (D-G) Glypican analysis in zebrafish. (D) Western blotting of ΔHS-stub using 3G10 antibody following heparinase III digestion of control and embryos injected with *COG4*^*WT*^ or *COG4*^*p.G516R*^ mRNA at 3 dpf. (E) Quantitation assay of glypican band density in (D) and two more replicates. The data are presented as mean ± SEM. Unpaired two-tailed *t*-test was used. *, p<0.05. (F) mRNA expression of glypican genes by RT-RCR using cDNA from control embryos at 6 hpf. (G) qPCR analyses of highly expressed glypicans in control and zebrafish embryos injected with human *COG4*^*WT*^ or *COG4*^*p.G516R*^ mRNA. Relative glypican level was normalized to β*-actin*. The graphs represent the 2^− ΔΔCt^ values. One-way ANOVA with Tukey’s multiple comparison tests was applied. ns, not significant; *, p<0.05, ***, p<0.001. Experiments were performed in triplicates with similar results.

We first checked glypican proteins using the same strategy as in SWS-derived cells. Interestingly, we found increased glypicans in embryos expressing *COG4*^*p.G516R*^ but not *COG4*^*WT*^ at 3 dpf (days post-fertilization) (Figure 2D-2E), consistent with the observation in SWS-derived fibroblasts. Zebrafish have ten glypicans expressing at different developmental stages (24). At 6 hpf (hours post-fertilization), we detected five glypicans as Gpc1b, 2, 4, 6a, and 6b by RT-PCR (Figure 2F), following with qPCR to compare their transcript level. We found that the *gpc2* and *gpc4* transcript levels are elevated in both *COG4*^*WT*^ and *COG4*^*p.G516R*^ embryos compared to uninjected control, but there is no significant difference between *COG4*^*WT*^ and *COG4*^*p.G516R*^ (Figure 2G). At 10 hpf, the transcript level of endogenous *gpc4* fell to a comparable level to control, with no distinction between *COG4*^*WT*^ and *COG4*^*p.G516R*^ (Figure 2G). *gpc2* mRNA level at 10 hpf was barely detectable.

### Expression of human *COG4*^*p.G516R*^ in zebrafish shortens body length and causes abnormal chondrocyte stacking and intercalation

We examined zebrafish body length at different stages to assess the developmental phenotypes caused by *COG4*^*p.G516R*^ expression. At the end of gastrulation, embryos expressing *COG4*^*p.G516R*^ variant exhibit shorter axis extension (Figure 3A), suggesting an abnormal convergent extension (CE) movement (17). Embryos expressing *COG4*^*WT*^ developed normally (Figure 3A). We tracked these injected embryos at later stages and found that expression of *COG4*^*p.G516R*^ causes shortened anterior-posterior (AP) body axis (Figure 3B) by an average of 18% at 3 dpf and 10% at 6 dpf (Figure 3C).

**Figure 3.**
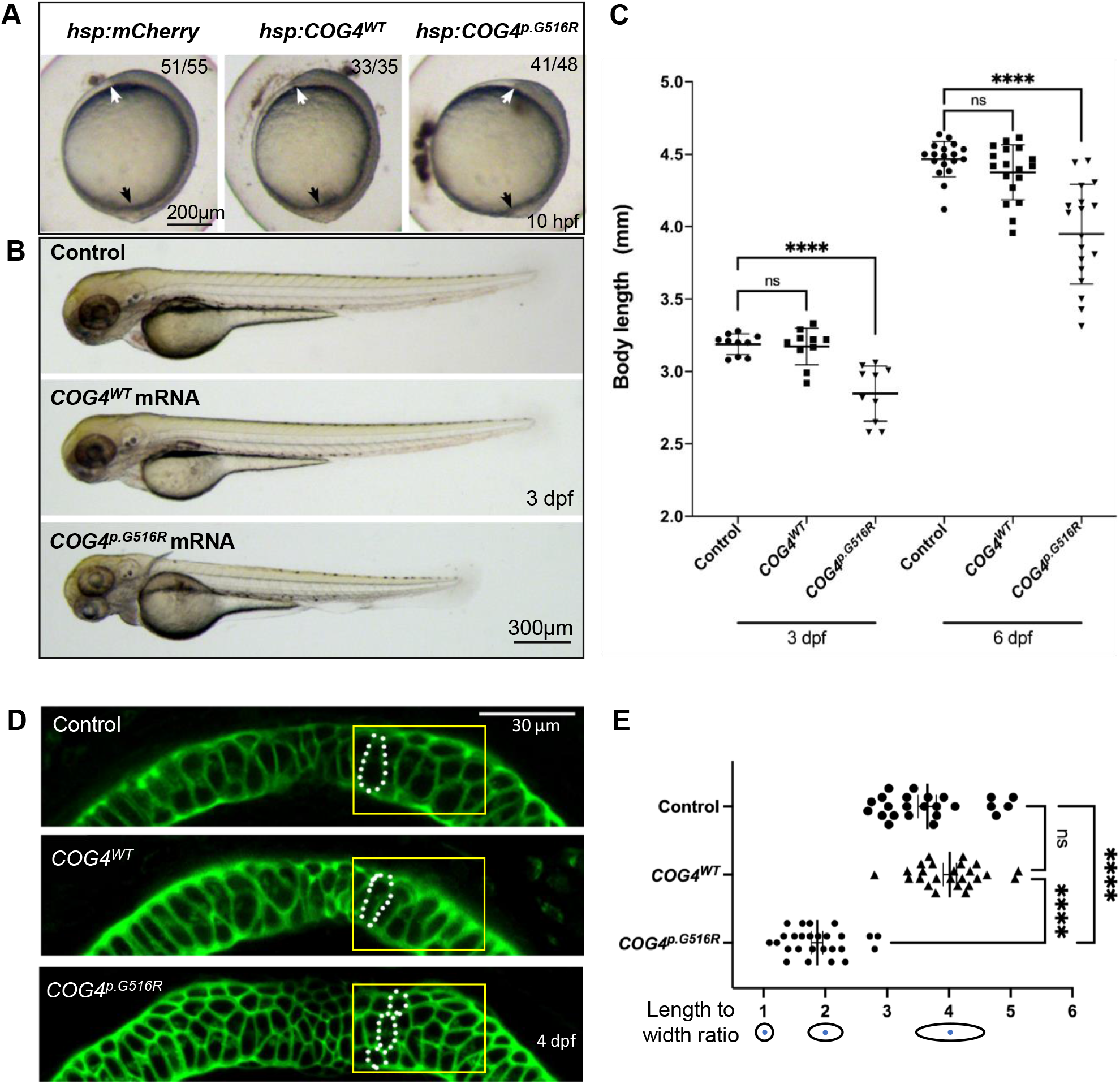
Expression of human *COG4*^*p.G516R*^ impairs zebrafish early development and chondrocyte intercalation. (A-B) Expression of *COG4*^*p.G156R*^ in zebrafish causes gastrulation defects and shortened body axis. (A) Lateral view of representative embryos. Anterior to the top. Compared to control construct *hsp70l:mCherry* and *COG4*^*WT*^, embryos expressing *COG4*^*p.G156R*^ show an axis extension defect at 10 hpf. White arrow points to the head region, and the black arrow points to the tailbud. Expression of *COG4*^*p.G156R*^ mRNA causes similar results. (C) Graphs show measured body length of each group at 3 dpf and 6 dpf. The data are presented as mean ± SD. One-way ANOVA with Tukey’s multiple comparison tests was applied. ****, p<0.0001; ns, not significant. (D) Expression of *COG4*^*p.G516G*^ causes craniofacial abnormalities. Ventral view of representative Meckel’s cartilage of zebrafish larvae at 4 dpf after WGA staining and imaged by confocal microscope. Dotted circular lines highlight chondrocyte cell shape and their relative configuration with each other. (E) Graphical representation of the length to width ratio of chondrocytes in the region of interest, yellow box in (D). Individual cell length/width ratio was measured in three representative Meckel’s cartilage images of each group. The data are presented as mean ± SEM. One-way ANOVA with Tukey’s multiple comparison tests was applied. ****, p<0.0001; ns, not significant. Experiments were performed in triplicates with similar results.

To investigate whether the presence of COG4^p.G516R^ in zebrafish impacts chondrocyte development, zebrafish at 4 dpf were stained with Wheat germ agglutinin (WGA), a lectin binding to glycoproteins in cartilage ECM to visualize chondrocyte morphology. In embryos expressing *COG4*^*p.G516R*^, Meckel’s cartilage was deformed, shown as defects in chondrocytes stacking and elongation (Figure 3D). Compared to *COG4*^*WT*^ embryos, more rounded chondrocytes were present in Meckel’s cartilage (Figure 3E). These defects can be observed as late as 7 dpf, as shown by Alcian blue staining (Figure S2). Chondrocyte stacking and intercalation problems further confirmed that abnormal glypican level could be one of the pathogenetic mechanisms involving in SWS. Other phenotypes seen in *COG4*^*p.G516R*^ injected embryos also include abnormal, stunted fin and cyclopia (Figure S3A-B).

### Expression of human *COG4*^*p.G516R*^ elevates *wnt4* transcript in zebrafish

Glypican 4 plays an essential role in gastrulation movements and contributes to craniofacial morphogenesis probably through Planar cell polarity (PCP)/non-canonical Wnt signaling (17, 25). Therefore, we assessed the expression of non-canonical *wnt*s by qPCR, and their spatial-temporal transcription pattern by whole-mount in situ hybridization. We found that *COG4*^*p.G516R*^ mRNA injected embryos contain more *wnt4* compared to controls (Figure 4A). In contrast, no significant changes were detected in a few other non-canonical *wnt*s, such as *wnt5b* and *wnt11f2*. Also, the upregulation of *wnt4* shows a dose-dependent response to the amount of *COG4*^*p.G516R*^ mRNA injected (Figure S4). This elevated transcript level of *wnt4* was further confirmed by whole-mount in situ hybridization (Figure 4B). At 6 hpf, *wnt4* was not detectable in either control or *COG4*^*WT*^ mRNA injected embryos, but it was in *COG4*^*p.G516R*^ mRNA injected embryos. At 10 hpf, *wnt4* is confined to the hindbrain in control and *COG4*^*WT*^ embryos, but *COG4*^*p.G516R*^ embryos had both increased and expanded expression of *wnt4. wnt11f2* transcript level did not change, but the distribution pattern was significantly altered. In uninjected control and *COG4*^*WT*^ embryos, *wnt11f2* was restricted to the dorsal midline. However, its expression pattern is dispersed in *COG4*^*p.G516R*^ embryos. No measurable difference was found for β-catenin protein, a marker of canonical Wnt signaling between *COG4*^*p.G516R*^ and controls (data not shown).

**Figure 4.**
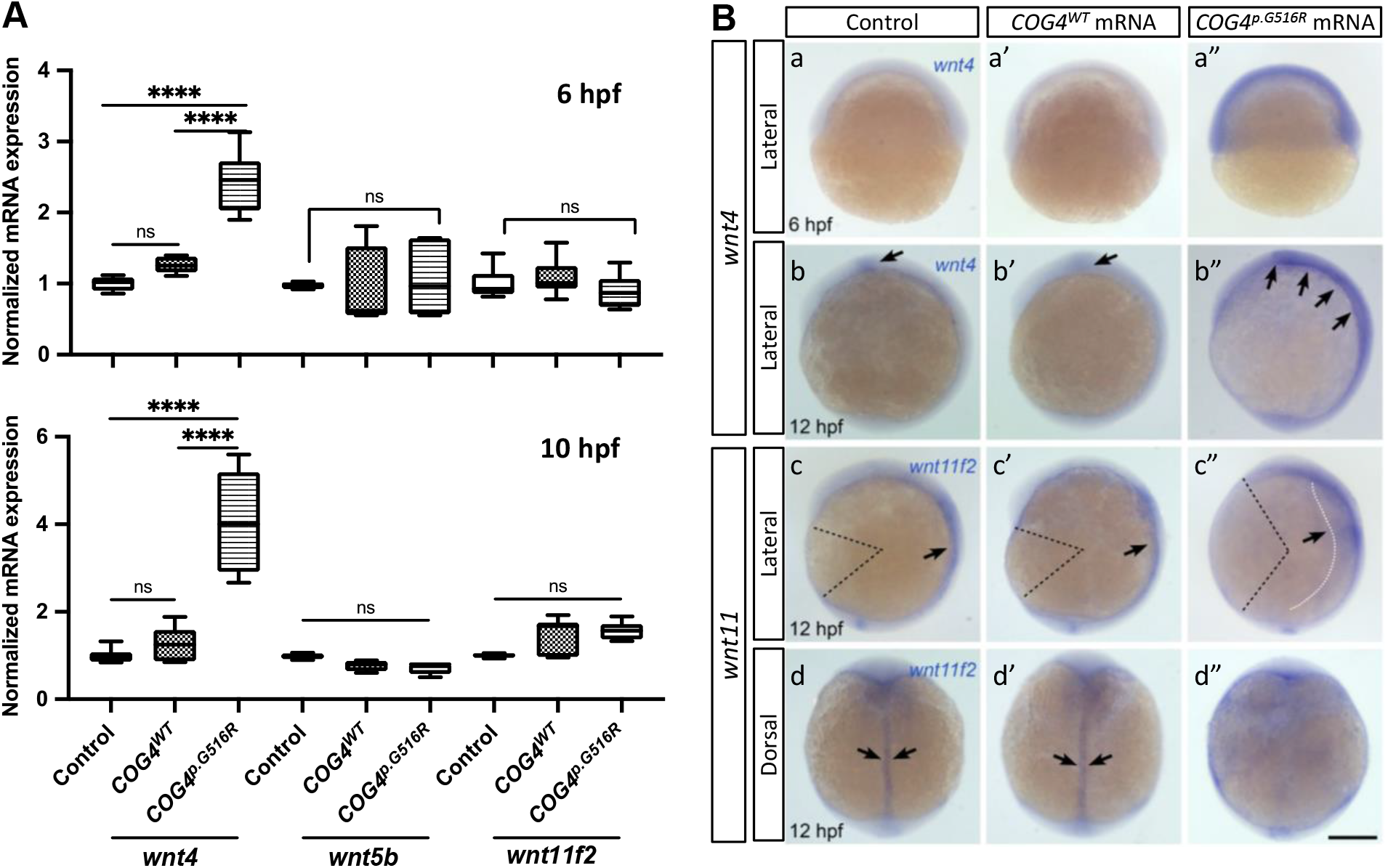
Expression of human *COG4*^*p.G516R*^ elevates *wnt4* transcript in zebrafish embryos. (A) Quantitative PCR analyses of selective non-canonical Wnt pathway ligands in zebrafish embryos injected with human *COG4*^*WT*^ or *COG4*^*p.G516R*^ mRNA. At 6 hpf, *wnt4* expression is significantly upregulated in embryos injected with *COG4*^*p.G516R*^ mRNA, but not with *COG4*^*WT*^ mRNA. Similar upregulation of *wnt4* transcripts is observed at 10 hpf as well. Bar graphs represent average gene expression relative to the housekeeping gene *β-actin*. The data are presented as mean ± SD. One-way ANOVA with Tukey’s multiple comparison tests was used. ****, p<0.0001; ns, not significant. (B) Representative images of whole-mount in situ hybridization from control and treated embryos. Whole-mount in situ hybridization analyses demonstrates that *wnt4* expression is elevated in embryos injected with *COG4*^*p.G516R*^ mRNA (a”, at 6 hpf; b’’ at 12 hpf), but there are no obvious changes in embryos injected with human *COG4*^*WT*^ mRNA (a’ and b’). (b, b’) Arrows point to the restricted *wnt4* expression domain in the hindbrain of zebrafish embryos at 12 hpf, (b”) arrows point to expanded expression of *wnt4* in the hindbrain region. At 12 hpf, *wnt11f2* expression intensity is not significantly changed in embryos injected with *COG4*^*p.G516R*^ mRNA (c”, d’’), compared to either control siblings (c, d) or human *COG4*^*WT*^ mRNA injected embryos (c’, d’). The dotted black color lines landmark the degree of angle between the head and tail, which is much more increased in embryos injected with *COG4*^*p.G516R*^ mRNA, suggesting defects in extension movement during gastrulation. Arrows in (c, c’ and d, d”) indicate that *wnt11f2* is restricted to the dorsal midline; however, its expression pattern is dispersed in *COG4*^*p.G516R*^ mRNA injected embryos, suggesting that the convergence movement is impaired (c”, white dotted line and arrow; d”, there is no clear midline expression). Scale bars: 200 µm. Experiments were performed in triplicates with similar results.

### Overexpression of zebrafish *wnt4* causes shortened body length and malformed Meckel’s cartilage

In zebrafish, *wnt4* overexpression has been studied at earlier embryonic stages and was found to inhibit cell movements without altering cell fates (17, 26). Thus, we evaluated the phenotypes of *wnt4* overexpression at a later embryonic stage. At 4 dpf, *wnt4* mRNA injected embryos showed shortened body length in response to the dosage of *wnt4* injected (Figure 5A). Interestingly, we also found abnormal chondrocyte stacking in Meckel’s cartilage (Figure 5B) and cyclopia (Figure 5C). These data show that overexpression of *wnt4* phenocopies shortened body length and malformed chondrocyte intercalation in *COG4*^*p.G516R*^ injected zebrafish.

**Figure 5.**
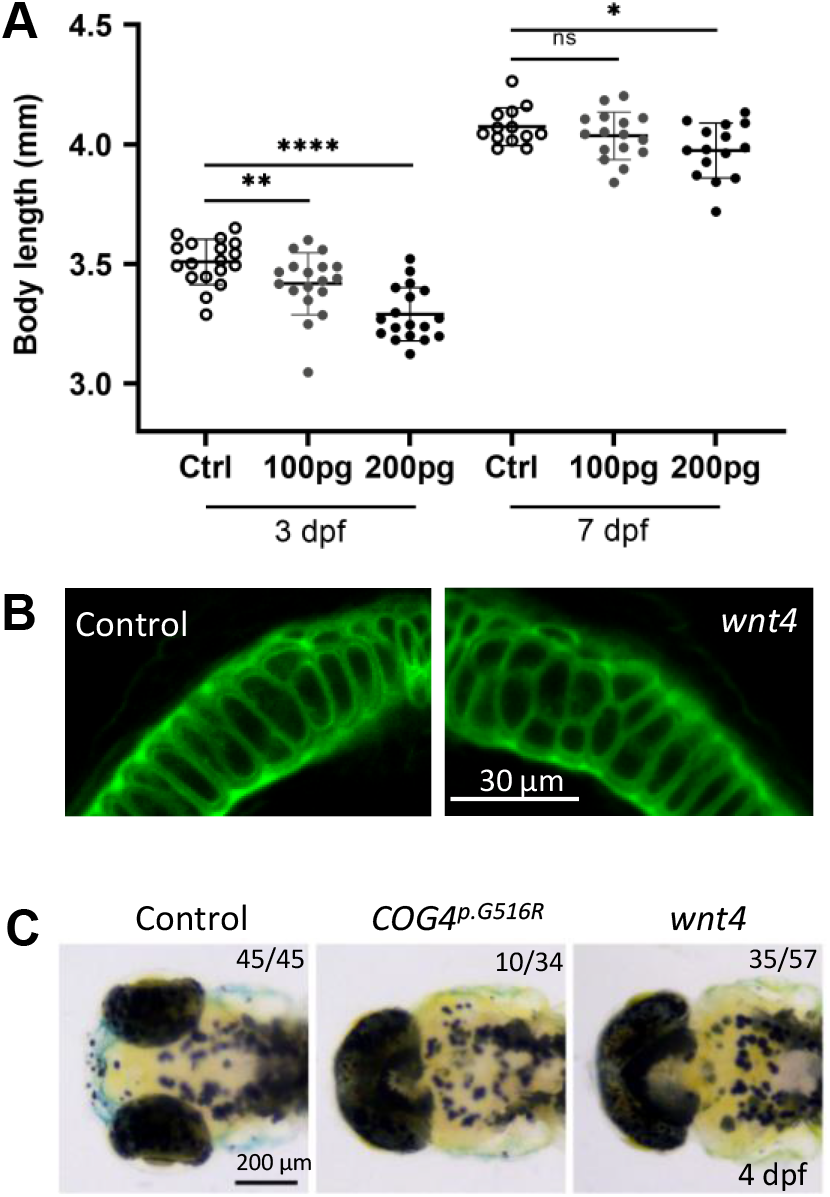
Overexpression of *wnt4* phenocopies zebrafish embryos injected with the *COG4*^*p.G516G*^ mRNA. (A) Graphs show measured body length of each group after *wnt4* mRNA injection at 3 dpf and 7 dpf. The data are presented as mean ± SD. Unpaired two-tailed *t*-test was used. ****, p<0.0001; **, p<0.01, *, p<0.05; ns, not significant. (B) Overexpression of *wnt4* causes abnormal chondrocytes stacking and intercalation at Meckel’s cartilage. Ventral view of representative Meckel’s cartilage of zebrafish larvae after *wnt4* injection at 4 dpf following WGA staining and imaged by confocal microscope. (C) Overexpression of *wnt4* causes cyclopia as expression of *COG4*^*p.G516G*^ variant. Dorsal view of representative images of cyclopia compared to control. 200pg of *wnt4* or *COG4*^*p.G516G*^ mRNA was used per embryo. Experiments were performed in triplicates with similar results.

### WNT inhibitor LGK974 suppresses the defects caused by *COG4*^*p.G516R*^ expression

We hypothesize the increased abundance of *wnt4* contributes to the pathogenetic mechanism for SWS. Therefore, reducing Wnt4 activity may suppress the developmental defects caused by the presence of COG4^p.G516R^. LGK974 is a pharmacological inhibitor of WNT porcupine O-acyltransferase (PORCN) which affects palmitoylation and secretion of Wnts. After optimizing the treatment procedure (Figure 6A), we found that 0.05 to 0.2 µM of LGK974 significantly shortened body length of control siblings (Figure S5) without causing optic cup morphogenesis and shorter tail induced by high concentrations of LGK974 (27). A 24-hour incubation with low concentrations (0.05 µM and 0.1 µM) of LGK974 restored shortened body length caused by *COG4*^*p.G516R*^ expression at 4 dpf (Figure 6B). Higher concentrations (0.15 and 0.2 µM) of LGK974 further shortened body length in *COG4*^*p.G516R*^ injected zebrafish, showing that there is a narrow optimal range for Wnt activity. This data suggests the imbalance of Wnt signaling may contribute to SWS pathogenesis. We also examined the chondrocyte morphology in Meckel’s cartilage and found that 0.05 and 0.1 µM of LGK974 is sufficient to suppress the deformed cartilage caused by *COG4*^*p.G516R*^ expression. Up to 0.1 µM LGK974 does not cause significant defects in control chondrocytes (Figure 6C-D).

**Figure 6.**
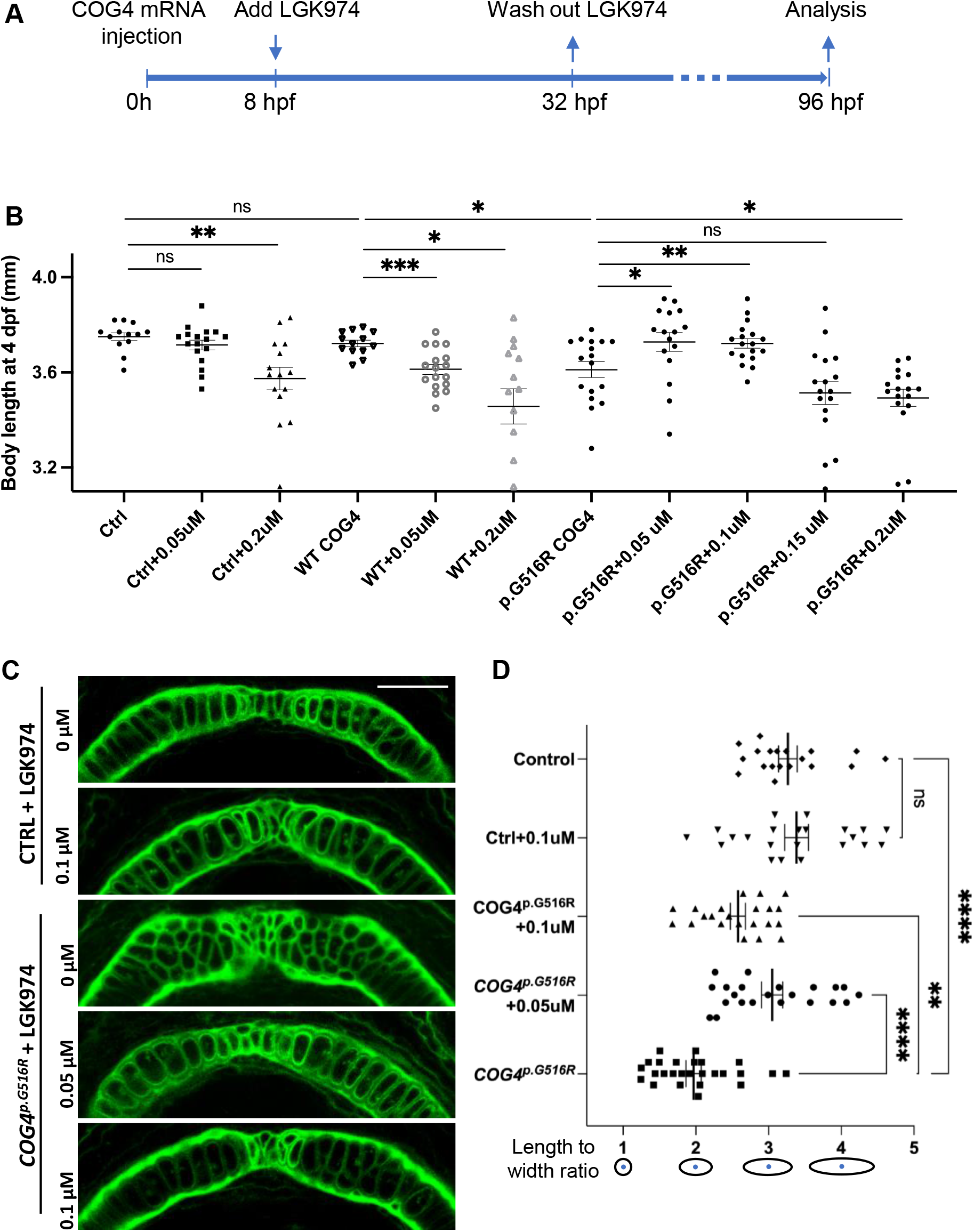
LGK974 treatment suppresses shortened body length and chondrocyte defect caused by of *COG4*^*p.G516G*^ expression in zebrafish. (A) Scheme of LGK974 treatment procedure. (B) Graphs show measured body length of each group at 4 dpf. The data are presented as mean ± SEM. Unpaired two-tailed *t*-test was used. ***, p<0.001; **, p<0.01; *, p<0.05; ns, not significant. (C) Ventral view of representative Meckel’s cartilage of control and *COG4*^*p.G516G*^ injected embryos with or without LGK974 treatment at 4 dpf following WGA staining and imaged by confocal microscope. (D) Graphs show the length to width ratio of chondrocytes in Meckel’s cartilage in (C) and two more representative Meckel’s cartilage images of each group. The data are presented as mean ± SEM. One-way ANOVA with Tukey’s multiple comparison tests was applied. ****, p<0.0001; **, p<0.01; ns, not significant. Experiments were performed in two biological replicates with similar results.

### WNT4 is elevated at mRNA and protein level in SWS-derived fibroblasts

To assess whether a similar mechanism was at play in human cells, we assayed transcript abundance of *WNT4* and other non-canonical *WNT*s in SWS individuals’ fibroblasts. Interestingly, both *WNT4* transcript and protein levels were increased in SWS individual fibroblasts (Figure 7A-B). Neither *WNT5a, WNT5b* and *WNT11* showed consistent significant differences in three SWS-derived cell lines (Figure 7A). We further detected downstream gene expression in the non-canonical pathway and found increased JNK phosphorylation (pJNK) in SWS cells compared to controls (Figure 7B), indicating an elevated non-canonical Wnt signaling. β-catenin, a marker for the canonical Wnt pathway, did not change (Figure 7B).

**Figure 7.**
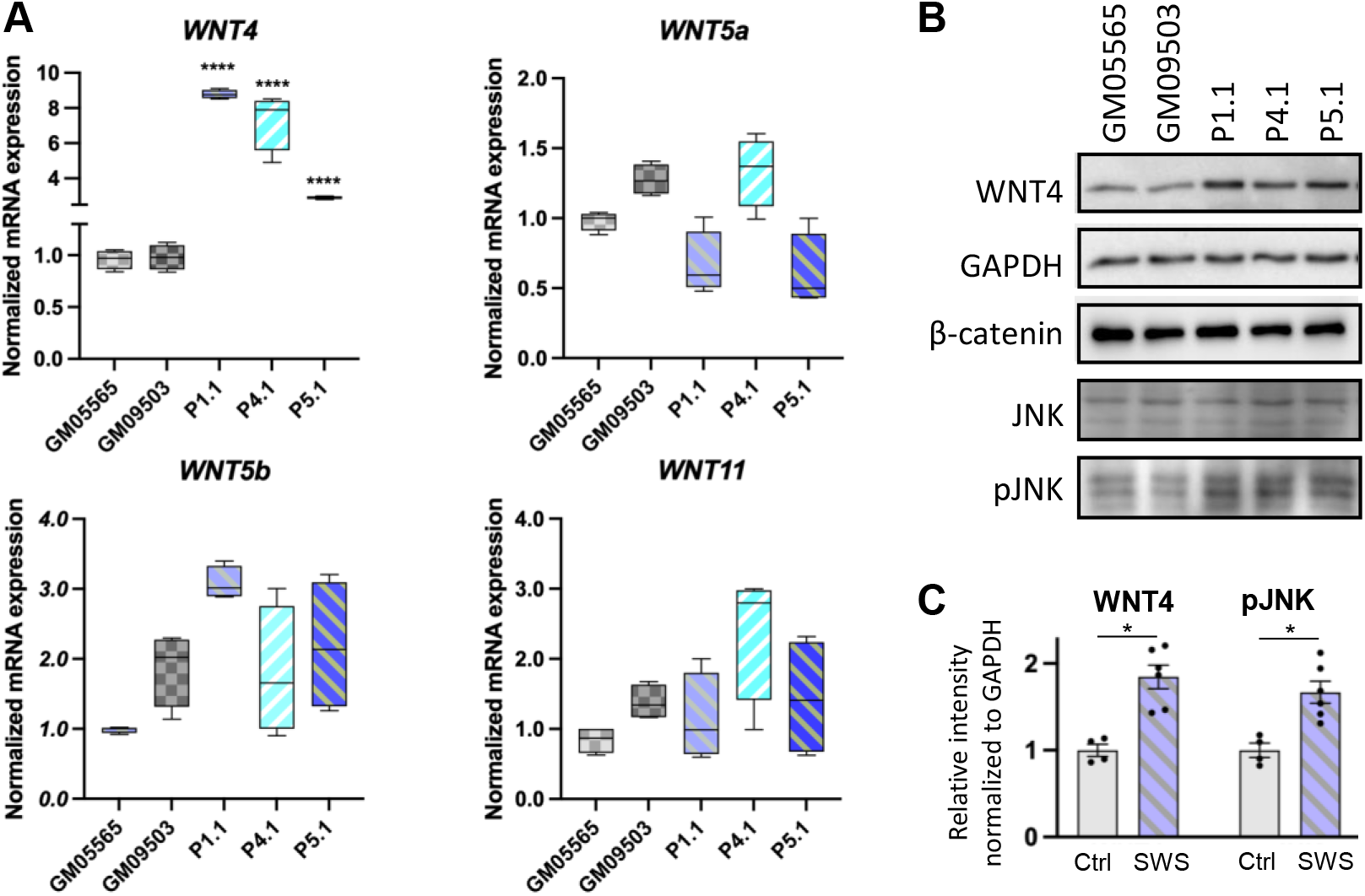
Non-canonical WNTs and related components level in SWS-derived cells. (A) qPCR analyses of selective non-canonical Wnt pathway components in SWS-derived fibroblasts. GAPDG was used as an internal control. The graphs represent the 2^−ΔΔCt^ values. Unpaired two-tailed *t*-test was applied for comparison of each SWS fibroblast with two controls. ****, p<0.0001. Experiments were performed in triplicates with similar results. (B) Western blotting of a few Wnt pathway components. GM05565 and GM09503 are control fibroblasts. P1.1, P4.1 and P5.1 are SWS-derived fibroblasts. (C) Quantitation assay of WNT4 and pJNK band density in (B) and the other replicates. The data are presented as mean ± SEM. Unpaired two-tailed *t*-test was used. *, p<0.05.

## Discussion

In this paper, we utilized SWS-derived fibroblasts and zebrafish, a vertebrate model to study SWS. In SWS-derived cells, the specific p.G516R amino acid substitution in COG4 selectively affected proteoglycans instead of global glycosylation. Increased glypicans were seen in both SWS-derived cells and *COG4*^*p.G516R*^ injected zebrafish embryos. This increase is probably due to impaired protein trafficking caused by COG4^p.G516R^ rather than changes in glypican transcript abundance. Glypicans are glycosylphosphatidylinositol-anchored proteins (GPI-APs) that consist of a conserved core glycan, phosphatidylinositol, glycan side chains and a protein moiety (28). In previous *gpc4* zebrafish studies, Topczewski demonstrated that Gpc4 regulates cellular movements during gastrulation by potentiating Wnt11f2 signaling and chondrocyte behavior independent of core Wnt pathway molecules (17, 25). Interestingly, low expression of *gpc4* suppressed Wnt11f2 defects, but high overexpression of *gpc4* inhibited the rescue, indicating a requirement for fine balance between Gpc4 and Wnt11f2 to ensure normal development. This is also true for COG4 function, since more than two-fold accumulation of COG4 leads to similar protein trafficking defects as seen in COG4-deficient cells (23). Therefore, *COG4*^*WT*^ is an essential control in this zebrafish study to confirm the expression of *COG4* is not excessive to impair its normal function.

Multiple studies have shown the role of glypicans in regulating Wnt signaling. In zebrafish, Gpc4 has been reported as a positive modulator of non-canonical Wnt signaling during zebrafish gastrulation (17). Studies in both *Drosophila* and *Xenopus* found glypicans can modulate the distribution of Wnt through binding the palmitoleate on Wnts and thus contribute to the release of signaling-competent Wnt (29-31). Among two groups of glypicans, Dally and Dally-like (Dlp) subfamilies, only Dlp has this palmitoleate-binding activity, which mainly includes Gpc4 and Gpc6 (29). Studies in chick embryos demonstrated Gpc4 in the neural crest enhances Wnt1/3a signaling and *wnt11* expression on the dorsomedial lip (32). Our data also suggests the role of glypicans in regulating Wnt signaling. We hypothesize that the accumulation of glypican(s) could induce the transcriptional upregulation of *wnt4*. However, lacking a specific Gpc4 antibody in zebrafish makes it difficult to confirm whether Gpc4 is the major player in causing these phenotypes seen in *COG4*^*p.G516R*^ injected embryos. Simply overexpressing *gpc4* in zebrafish seems to show no significant defects (16). We speculate the abnormalities seen in *COG4*^*p.G516R*^ injected embryos are due to disrupted glypican trafficking rather than changes in glypican expression. Further studies on how glypican and wnt4 crosstalk is yet to be demonstrated. Whereas glypicans are the most impaired proteoglycans caused by the heterozygous variant in COG4 (p.G516R), we could not rule out other proteoglycans may also contribute to the pathogenesis of SWS. Although *dcn* and *cspg4* mutants in zebrafish do not show chondrocyte intercalation problems, they may cause shorter body axis or other types of cartilage malformations (11, 12).

Besides increased glypicans, it is interesting that we also found elevated *wnt4* transcript in SWS-derived cells and *COG4*^*p.G516R*^ injected zebrafish embryos. Wnts are a group of secreted glycoproteins involved in cell-cell signaling (33, 34). To date, 22 Wnts have been identified in vertebrates (The Wnt homepage: http://web.stanford.edu/group/nusselab/cgi-bin/wnt/). Among those, Wnt4 has shown important roles in embryonic development, skeletal/bone regeneration, sex determination with additional developmental defects in a cell-specific and tissue-specific manner (35-39). Most importantly, Wnt4 plays a pivotal role in regulating chondrocyte differentiation. In chick limb studies, *Wnt4* expression is found in joint cells and cartilage and its misexpression accelerates maturation of chondrocytes, causing shortened long bones (40-42). An *in vitro* study also found that Wnt4 blocks the initiation of chondrogenesis and accelerates terminal chondrocyte differentiation (41). Misexpression of Z*wnt4* (zebrafish *wnt4*) and *Xwnt4* (Xenopus *wnt4*) in zebrafish causes shortened trunk and tail (26) and abnormal chondrocyte intercalation in Meckel’s cartilage (our data), confirming Wnt4 is involved in regulation of chondrocyte behavior, like *gpc4* and *wnt5b* (25). In mice, Wnt4 defects led to short-limb dwarfism (43). Most relevant, conditional expression of *Wnt4* in mice also causes dwarfism with small skeletons, dome-shaped skull and small jaws (44). Both *Wnt4* defects and overexpression cause dwarfism, indicating Wnt4 functions within a narrow range for normal development, and bidirectional disturbance causes developmental abnormalities.

LGK974 is a PORCN inhibitor, that blocks PORCN-dependent palmitoylation and secretion of WNT family ligands (45). Currently, it is being evaluated as an anticancer agent against a broad range of diseases associated with deviant Wnt signaling (45, 46). LGK974 has been used to inhibit canonical and non-canonical signaling in multiple studies as a WNT modifier (47-49). In our study, titrating LGK974 allowed us to regulate the amount of Wnt during development. At low concentrations, it suppressed and nearly normalized the shortened body length caused by the presence of COG4^p.G516R^, probably by inhibiting Wnt4 (or other Wnts) palmitoylation and secretion. This rescue experiment not only supports altered Wnt signaling as one of the possible pathogenesis mechanisms contributing to SWS, but also suggests a potential therapeutic strategy for SWS individuals.

In our SWS zebrafish model, the specific heterozygous dominant variant in COG4 (p.G516R) shows elevated glypicans and *wnt4*, along with developmental defects. Compared to COG4 KO zebrafish (3, 20), it is impressive that the single mutation in COG4 is sufficient for shortened body length, and abnormal chondrocyte stacking. However, there are still some differences between COG4 KO and *COG4*^*p.G516R*^ zebrafish; for example, the single variant in COG4 doesn’t decrease the overall GAG amount shown in COG4 KO zebrafish by Alcian blue staining. *COG4*^*p.G516R*^ larvae also show cyclopia and distinct pectoral fins phenotype compared to COG4 KO zebrafish.

In summary, using SWS-derived fibroblasts and a zebrafish model, we demonstrated that the specific dominant variant COG4^p.G516R^ causes the accumulation of GPI-anchored glypicans, which most likely involves COG4-dependent altered trafficking of these proteins. In zebrafish, the presence of COG4^p.G516R^ also elevates *wnt4* transcript and causes chondrocyte morphogenesis defects, which might explain the short stature and distinctive craniofacial features in SWS individuals. WNT4 and non-canonical Wnt signaling component pJNK are also elevated in cultured SWS-derived fibroblasts. We further demonstrate the Wnt inhibitor LGK974 could suppress the defects caused by *COG4*^*p.G516R*^ in zebrafish. These results from SWS-derived cell lines and zebrafish point to altered non-canonical Wnt signaling as one possible mechanism underlying SWS pathology. How increased COG4^p.G516R^ leads to elevated *WNT4* in SWS cells and zebrafish is still unknown. Transgenic and CRISPR knock-in SWS zebrafish lines are under development to address these issues.

## Materials and Methods

### Cell cultures

Dermal primary fibroblasts derived from healthy controls (GM08429, GM08680, GM03349, GM05565 and GM09503) were obtained from Coriell Institute for Medical Research (Camden, NJ). Each SWS-derived fibroblast line was obtained by the referring clinician and grown via a clinical lab service and then sent to us with consent through an approved IRB.

Fibroblasts were cultured in Dulbecco’s Modified Eagle’s medium (DMEM) containing 1 g/L glucose supplemented with 10% heat-inactivated fetal bovine serum (FBS) and 1% antibiotic-antimycotic (Life Technologies, Carlsbad, CA, USA).

### Zebrafish husbandry

All zebrafish experiments were performed in accordance with protocols approved by SBP IACUC. Zebrafish were maintained under standard laboratory conditions at 28.5 °C. Embryos were staged according to Kimmel *et al*. (50).

### Immunoblotting

For glypicans analysis, fibroblasts were treated with 5mU/ml Heparinase III in serum-free medium for 1 hour at 37□C. Zebrafish larvae were homogenized and treated with 25mU Heparinase III in H buffer (20 mM Tris-HCl, pH 7.0, 0.1 mg/ml BSA, and 4 mM CaCl_2_) for 1.5 hours at 37□C. Samples were harvested using SDS lysis buffer (62.5 mM Tris-HCl, pH 6.8, 2% SDS, and 10% glycerol) supplemented with protease and phosphatase inhibitors (Sigma-Aldrich) as previously described (3). For analysis of PCP pathway components, 2×10^4^ fibroblast cells were seeded to 6-well plates and harvested after two days using SDS lysis buffer. Equal amounts of denatured proteins were separated via SDS–polyacrylamide gel electrophoresis following transfer and antibody inoculation as described previously (51). Antibodies used were Δ-Heparan Sulfate (AMSBIO, F69-3G10), Chondroitin 6 Sulfate (Millipore, MAB2035), GAPDH (Invitrogen, MA5-15738), WNT4 (R&D, MAB4751), mCherry (Rockland, 600-401-P16), COG4 (provided by Dr. Daniel Ungar, University of York, UK), β-catenin (Santa Cruz, sc-7963), JNK (sc-7345), pJNK (sc-293136).

### mRNA expression analysis

Total RNA was extracted from cells or zebrafish embryos using TRIzol™ (ThermoFisher, 15596081) reagent according to manufacturer protocol. cDNA was synthesized using QuantiTect Reverse Transcription Kit (QIAGEN, 205311). qPCR and data analysis were performed as described previously (51). Briefly, primer pairs targeting genes of interest were designed using NCBI Primer-BLAST and available upon request. qPCR reactions were performed with PowerUp SYBR® Green PCR Master Mix (ThermoFisher, A25742). The standardized cycle conditions were applied in Applied Biosystems 7900HT Fast Real-Time PCR System. SDS2.3 software was used to analyze expression data of reference genes. The mRNA levels were normalized to the levels of housekeeping genes, *GAPDH* for fibroblasts and β*-actin* for zebrafish, and 2^− ΔΔCt^ values were calculated and compared.

### Immunofluorescence

Whole-mount immunofluorescence was performed as previously described (52), using lectin WGA (Vector Laboratories, FL1021) against cell membrane. Fluorescence images were acquired using an LSM 510 confocal microscope (Zeiss, Germany) with a 40x water objective. Digital images were processed with Adobe Creative Suites.

### Cloning of human *COG4* and in vitro mRNA synthesis

Full length human *COG4* was cloned into pCS2+ vector from plasmid hCOG4-siR-3myc in AAZ6 (Gift from Professor Vladimir V. Lupashin) using In-Fusion® HD Cloning Kit (TaKaRa Bio, 638909) with primers 5’-ATGGGAACCAAGATGGCGGA-3’, 5’-TTACAGGCGCAGCCTCTTGATAT-3’, 5’-ATCAAGAGGCTGCGCCTGTAATCAAGGCCTCTCGAGCCTCT-3’ and 5’-TCCGCCATCTTGGTTCCCATATTCGAATCGATGGGATCCT-3’. SWS point mutation of G to A was generated using Q5® Site-Directed Mutagenesis Kit (NEB, E0554S) with primers 5’-CATCCAGCGCaGGGTGACAAG-3’ and 5’-TCCTGGAAGGTGGTGGCA-3’. WT *COG4* and SWS *COG4* mRNAs (*COG4*^*WT*^ and *COG4*^*p.G516R*^) were synthesized using Invitrogen mMESSAGE mMACHINE SP6 Transcription Kit (Thermo Fisher, AM1340) following restriction enzyme NotI digestion and purified using MEGAclear(tm) Transcription Clean-Up Kit (Thermo Fisher, AM1908). 100pg mRNA was injected to each zebrafish embryos unless state otherwise. As a complementing strategy, *Hsp70l: COG4*-*P2A-mCherry; cryaa:dsRED and* SWS *COG4* were generated by cloning coding sequence for WT *COG4* and SWS *COG4* downstream of the Hsp70l promoter in the parent vector *hsp70l:-LateSV40pA; cryaa:dsRED-RabBGpA; I-Sce* vector (Gift from Dr. Joseph Lancman). I-SceI meganuclease (NEB, R0694) was co-injected with 20pg of the constructs per embryo.

### Cloning of full-length zebrafish *wnt4* and synthesizing antisense probes

Full length zebrafish *wnt4* was cloned from cDNA into pCS2+ vector using In-Fusion® HD Cloning Kit with primers: 5’-CCCATCGATTCGAATATGTCATCGGAGTATTTGATAAGGT-3’, 5’-CTCGAGAGGCCTTGATCACCGACACGTGTGCAT-3’, 5’-TCAAGGCCTCTCGAGCCTCT-3’ and 5’-ATTCGAATCGATGGGATCCTGCA-3’. *wnt4* mRNA synthesis and purification were performed as described above. 100pg mRNA was injected into each zebrafish embryos unless mentioned otherwise. For antisense RNA probe synthesis, the full length of *wnt4* including 3’-UTR was cloned into pGEM-T easy vector (Promega, A1360) by primers 5’-ATGTCATCGGAGTATTTGATAAGG-3’ and 5’-AGTCTTTGACACAGCATATATTTC-3’ from cDNA. After verifying insertion direction by sequencing, the antisense RNA probe was then synthesized with SP6 RNA polymerase following ApaI digestion.

### Standard whole-mount in situ hybridization

Standard whole-mount in situ hybridization was performed as described previously (53). INT/BCIP (175 μg/ml; Roche) was used as alkaline phosphatase substrates. The following molecular markers were used: *wnt4*, and *wnt11f2* (Gift from Dr. Diane Sepich, previously used in (54, 55)).

### Wnt inhibition assay

LGK974 (Cayman Chemical, No. 14072) was dissolved in DMSO to make 10 µM stock. Different concentrations of LGK974 were added to control or injected zebrafish groups at 8 hpf for 24 hours. 0.01% DMSO was used as a vehicle.

## Ethics

This study was performed in strict accordance with the recommendations in the Guide for the Care and Use of Laboratory Animals of the National Institutes of Health. All of the animals were handled according to approved institutional animal care and use committee (IACUC) protocols of SBP.

## Acknowledgements

This work was supported by The Rocket Fund and R01DK99551. Z-J.X is a Rocket Fund Fellow. The authors thank Dr. Heather Flanagan-Steet for providing reagents and guidance and for critical reading of the manuscript. The authors thank Dr. Yu Yamaguchi for critical discussion. The authors thank Dr. Diane Sepich for providing *wnt11f2* DNA construct for in situ probe synthesis.

---

**Figure S1. Western blots of CSPG following Chondroitinase ABC digestion to generate CS-stubs**. C1 and C2 are control fibroblasts. C1, GM0849; C2, GM0860. P1.1, P4.1 and P5.1 are SWS-derived fibroblasts.

**Figure S2. Expression of *COG4***^***p*.*G156R***^ **in zebrafish shows abnormal chondrocytes intercalation of Meckel’s cartilage at late developmental stage**. Ventral view of representative Meckel’s cartilage of zebrafish larvae at 7 dpf after Alcian Blue staining, imaged by light dissecting microscope. Dotted circular lines highlight cell shape and their relative configuration with each other.

**Figure S3. Expression of *COG4***^***p*.*G156R***^ **in zebrafish causes cyclopia, stunted fin, and abnormal ceratohyal cartilage in zebrafish**. (A) Expression of *COG4*^*p.G516G*^ causes cyclopia in zebrafish. Dorsal view of representative images of cyclopia compared to control and *COG4*^*WT*^. 200pg of *COG4*^*WT*^ or *COG4*^*p.G516G*^ mRNA was injected into each embryo. (B) Ventral view of representative zebrafish larvae at 6 dpf after Alcian Blue staining, imaged by light dissecting microscope. 20pg DNA was injected into each embryo, and heat shock was performed at 24 hpf and 46 hpf for 1 hour each at 38°C degree.

**Figure S4. *wnt4* upregulation shows a dose-dependent response to *COG4***^***p*.*G516R***^ **mRNA**. Bar graphs represent quantitative PCR analyses of *wnt4* at 6 hpf after injection of different amounts of *COG4*^*p.G516R*^ mRNA. *wnt4* expression was normalized to β*-actin*. ****, p<0.0001; **, p<0.01; ns, not significant.

**Figure S5. Dose-dependent response analysis of Wnt inhibitor LGK974 in zebrafish**. (A) Body length at 3dpf after LGK974 treatment. (B) Representative images after LKG974 treatment at 3dpf. The data are presented as mean. ****, p<0.0001.

## Notes

### Competing Interest Statement

The authors have declared no competing interest.

