## Supplementary figures and images for "A single heterozygous mutation in *COG4* disrupts zebrafish early development via Wnt signaling"

### Supplemental figures

Figure S1.

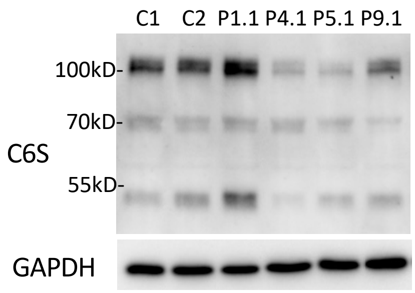

Figure S2

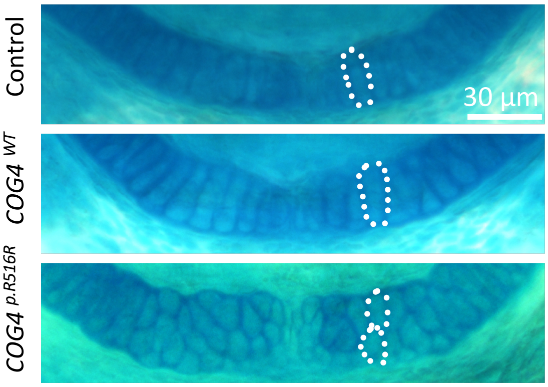

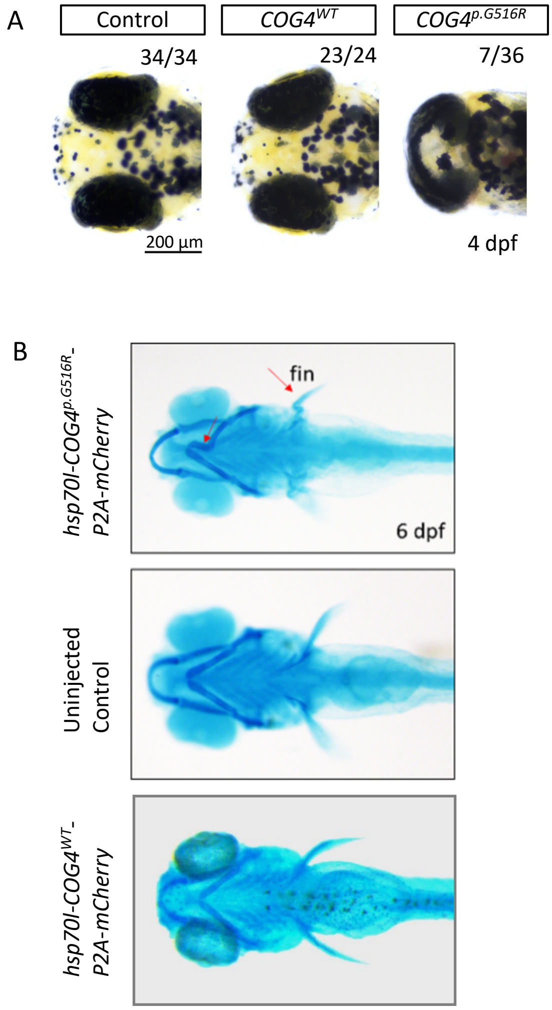

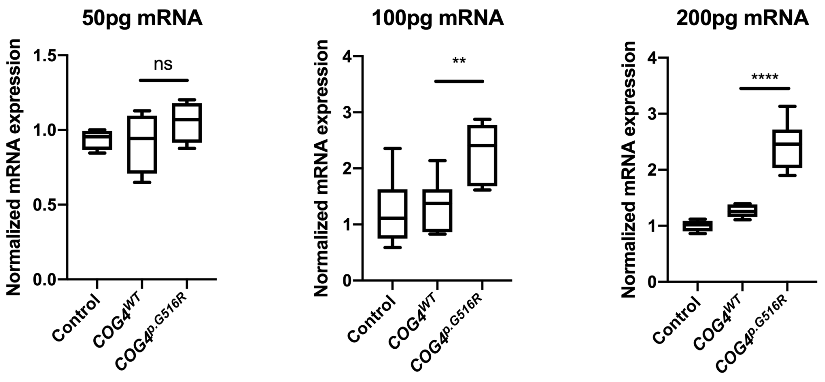

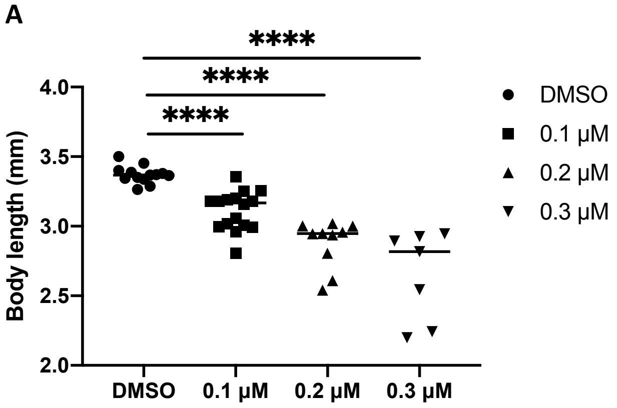

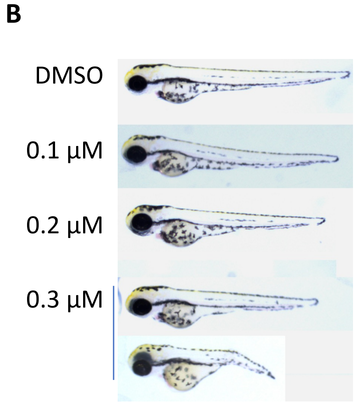
